# CluSim: a Python package for the comparison of clusterings and dendrograms

**DOI:** 10.1101/410084

**Authors:** Alexander J. Gates, Yong-Yeol Ahn

**Affiliations:** Center for Complex Networks Research, Northeastern University, Boston, 02115, USA; Department of Informatics, Indiana University, Bloomington, 47408, USA; Program in Cognitive Science, Indiana University, Bloomington, 47408, USA.

## Abstract

Quantifying the similarity of clusterings is a fundamental step in data analysis. Clustering similarity is the basis for method evaluation, consensus clustering, and tracking the temporal evolution of clusters, among many other tasks. Here we provide *CluSim*, a comprehensive Python package for the comparison of partitions, overlapping clusterings, and hierarchical clusterings (dendrograms) with more than 20 similarity measures. The *CluSim* package provides both analytic and empirical methods for assessing the similarity of clusterings in the context of a random model, and provides the novel element-centric approaches for clustering similarity measure that we introduced recently. We illustrate the use of the package through two examples: an evaluation of the clustering of Gene Expression data in the context of different random models, and detailed analysis of model incongruence using element-centric comparisons between a set of phylogentic trees (dendrograms).

**Availability and implementation:** The CluSim Python package and accompanying jupyter notebook is available at https://github.com/Hoosier-Clusters/clusim with the MIT open source licence.

**Contact:** **or**

## 1. Introduction

Clustering is a primary method to reveal the structure of data [1]. To understand, evaluate, and leverage data clusterings, we need to quantitatively compare them. Clustering comparison is the basis for method evaluation, consensus clustering, and tracking the temporal evolution of clusters, among many other tasks. For instance, clustering method evaluation is usually achieved by comparing the method’s result to a planted reference clustering, under the assumption that the more similar the method’s solution is to the reference clustering, the better the method. Despite the importance of clustering comparison, no consensus has been reached for a standardized assessment; each similarity measure rewards and penalizes different criteria, sometimes producing contradictory conclusions. Each scientific community has adopted their own standard practices, often without considering whether the measures’ underlying assumptions are appropriate for the given task.

Clustering similarity measures can be classified based on the cluster types: i) *partitions* that group elements into non-overlapping clusters, ii) *hierarchical clusterings* that group elements into a nested series of partitions (a.k.a. dendrogram), or iii) *overlapping clusterings* with elements belonging to multiple clusters. Furthermore, in order to establish a baseline and interpret the similarity score, it is often argued that clustering similarity should be assessed in the context of a random ensemble of clusterings. Such a correction procedure requires two choices: *a model for random clusterings* and *how clusterings are drawn from the random model*. With few exceptions, similarity measures are only designed to compare clusterings of the same type, and the decisions required for the correction procedure are usually ignored or relegated to the status of technical trivialities even though such decisions can sometimes reverse one’s conclusions completely [2].

Here, we introduce *CluSim*, a python package that provides a unified library of over 20 clustering similarity measures for partitions, dendrograms, and overlapping clusterings. To our knowledge, this package constitutes the first collection of clustering similarity measures for all three clustering types and extended access to random models of clusterings. The package also includes the element-centric similarity measure that naturally unifies the comparison of all three clustering types [3]. We illustrate the use of the package through two examples: comparing clusterings of Gene Expression data in the context of random models, and element-centric comparisons between a set of phylogentic trees (dendrograms).

## 2. Materials and Methods

The basic class in the *CluSim* package is a *Clustering*, or an assignment of labeled elements (i.e. data points or network vertices) into clusters (the groups). A *hierarchical Clustering* also contains a dendrogram, or more generally an acyclic graph, capturing the nested structure of the clusters. In *CluSim*, a *Clustering* can be instantiated from 7 common formats, including full support for *scipy, scikit-learn*, and *dendropy* clustering formats [4, 5, 6].

*CluSim* provides more than 20 clustering similarity and distance measures for the comparison between two *Clusterings*. All similarity measures produce a score in the range [0, 1], where 1 indicates identical clusterings and 0 indicates maximally dissimilar clusterings. See the online documentation for a detailed list and mathematical definitions of these similarity measures.

To facilitate comparisons within a set of clusterings, the *CluSim* package provides two implementations of the correction for chance. Analytic solutions are available for the Rand index and Normalized Mutual Information using five random models: the permutation model, both one-sided and two-sided models for clusterings with a fixed number of clusters, and both one-sided and two-sided models for all random clusterings [7, 8, 2]. For all other similarity measures, the correction for chance is estimated by randomly sampling the random ensemble of *Clusterings* using the provided random Clustering generators.

## 3. Applications

To illustrate the use of our package, we offer two examples that highlight insights gained from an advanced package on clustering similarity.

In the first example, we assess the performance of Hierarchical Clustering to identify cancer type from gene expression data. Specifically, we compare a ground truth classification of patient cancer type to a derived classification using agglomerative hierarchical clustering of gene expression [9] in the context of two different random models. The traditional approach assumes the Permutation Model for random clusterings in which the number and sizes of clusters are held fixed. As can be seen in Fig. 1, the true comparison (red line) is larger than the mean of pair-wise comparisons between clusterings in the permutation model (black line, blue histogram), suggesting the similarity is greater than expected by chance. However, a more appropriate random model for the given scenario is the one-sided model with a fixed number of clusters [2]. Fig. 1 (bottom) shows that the true comparison (red line) is actually smaller than the mean of pair-wise comparisons between clusterings in the random model (black line, blue histogram), suggesting that a randomly generated clustering would be more similar to the ground truth clustering than the computationally derived solution.

**Figure 1:**
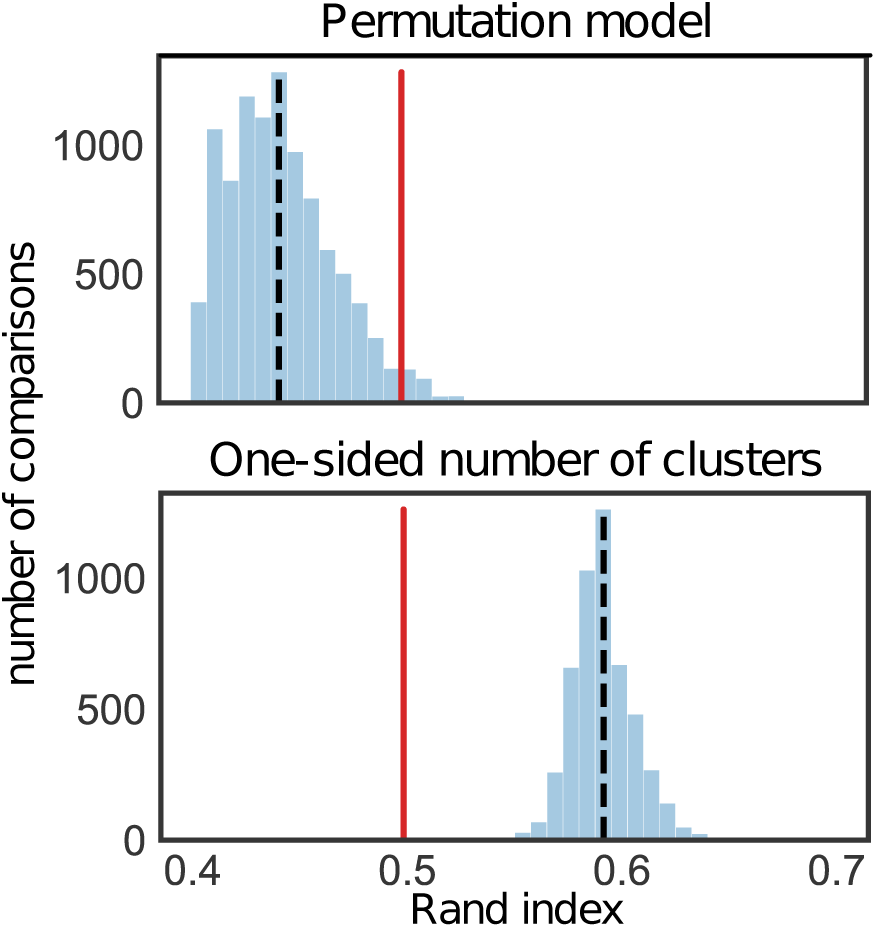
Evaluating clustering comparisons w.r.t. random models. A comparison using the Rand Index between the classification of cancer types and clustering labels derived using Hierarchical Clustering on gene expression data (0.5, red). above, Pairwise comparisons between samples from the Permutation model (blue, see Hubert & Arabie[7]) with mean 0.44 (black). below, Pairwise comparisons between samples from the one-sided model with a Fixed Number of Clusters (blue, see Gates & Ahn[2]) with mean 0.59 (black). The Permutation model suggests Hierarchical Clustering is more similar to the ground truth than a random clustering, while the one-sized fixed number of clusterings model, the more appropriate model for this scenario, reveals that the result is less similar than random clusterings.

In the second example, we identify the loci of gene-tree heterogeneity in an analysis of 424 nuclear genes from 37 eutherian mammals [10]. Specifically, we perform all pair-wise comparisons between the 424 phylogenetic trees (dendrograms, exemplified in Fig. 2a) derived from individual gene sequence data [11]. The average element-centric similarity between the trees (Fig. 2b) reveals their overall similarity, with few conflicts near the roots of the trees (high similarity for the scaling parameter, *r <* 0), while decreasing similarity suggest greater conflicts towards the leaves (lower similarity for the scaling parameter,*r >* 0). The distribution of element-wise frustration scores over the taxa reveal the loci of greatest gene tree in-congruence (Fig. 2c). Specifically, the 8 taxa with lowest frustration correspond to the 5 taxa previously identified with structural discrepancies (bats, shrews, and hedgehog, blue, [11]), and the 3 taxa with the smallest bootstrapping support in the maximum-pseudolikelihood coalescent tree (pig, guinea pig, kangaroo rat, purple, [10]). This comparison provides quantitative insights into the complexities commonly observed in phylogentic data.

**Figure 2:**
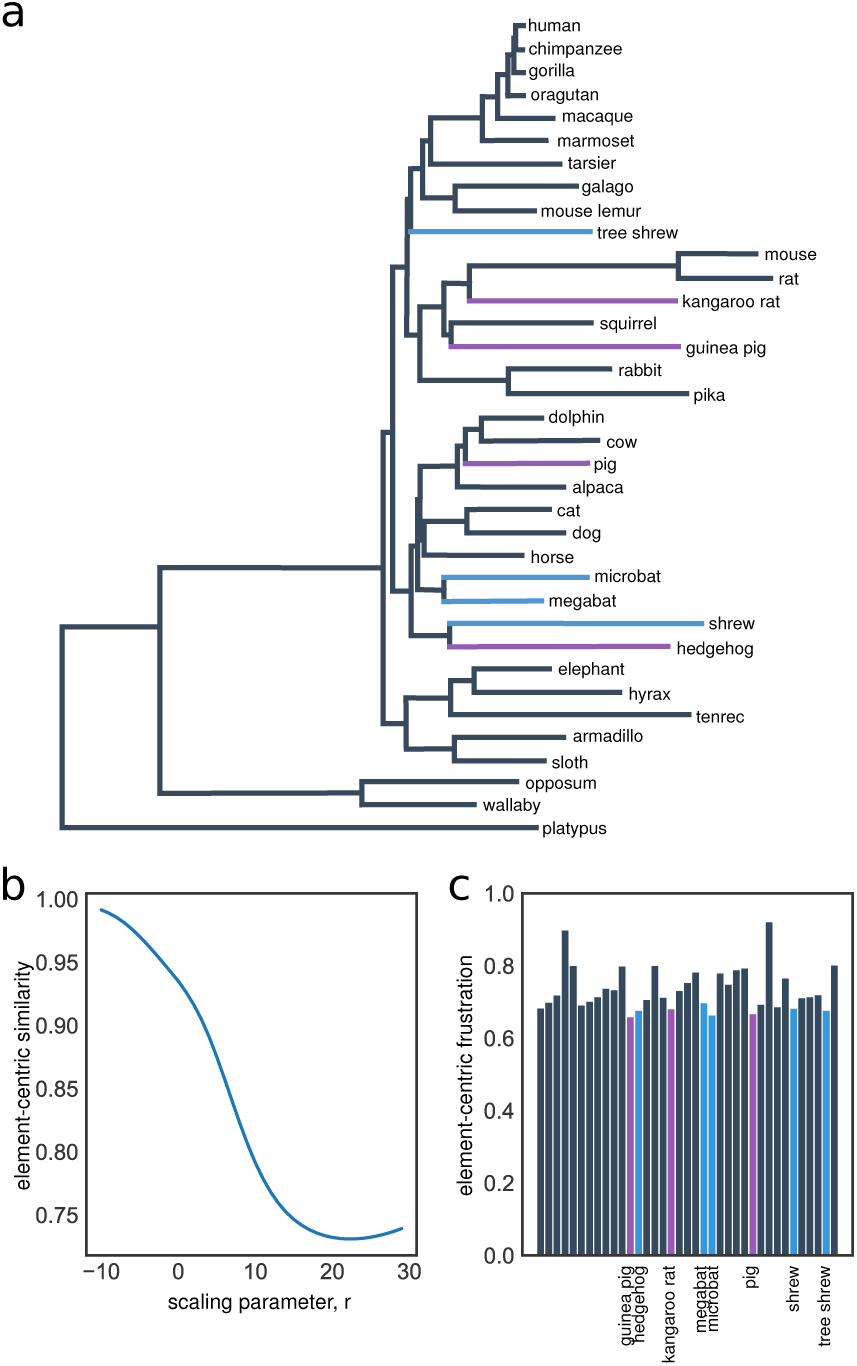
Element-centric comparisons of phylogenetic dendrograms. **a**, An example phylogenetic tree for the 37 mammals from [10]. **b**, The average element-centric similarity between 424-gene trees for different scaling parameters reveals few conflicts near the roots (left, *r <* 0), while decreasing similarity for increasing *r* suggests greater conflicts towards the leaves (left, *r >* 0). **c**, The element-centric frustration highlights the 5 taxa previously identified with structural discrepancies (bats, shrews, and hedgehog, blue, [11]), and the 3 taxa with the smallest bootstrapping support in the maximum-pseudolikelihood coalescent tree (pig, guinea pig, kangaroo rat, purple, [10]).

## Acknowledgements

The authors would like to thank Ian Wood for thoughtful discussions.

## References

[1] Anil K. Jain, M. Narasimha Murty, and Patrick J. Flynn. Data clustering: a review. ACM Computing Surveys (CSUR), 31(3):264–323, 1999.

[2] Alexander J. Gates and Yong-Yeol Ahn. The impact of random models on clustering similarity. Journal of Machine Learning Research, 18(87):1–28, 2017.

[3] Alexander J. Gates, Ian B. Wood, William P. Hetrick, and Yong-Yeol Ahn. On comparing clusterings: an element-centric framework unifies overlaps and hierarchy, 2018. arXiv:1706.06136.

[4] Eric Jones, Travis Oliphant, Pearu Peterson, et al. SciPy: Open source scientific tools for Python, 2001–.

[5] F. Pedregosa, G. Varoquaux, A. Gramfort, V. Michel, B. Thirion, O. Grisel, M. Blondel, P. Prettenhofer, R. Weiss, V. Dubourg, J. Vanderplas, A. Passos, D. Cournapeau, M. Brucher, M. Perrot, and E. Duchesnay. Scikit-learn: Machine learning in Python. Journal of Machine Learning Research, 12:2825–2830, 2011.

[6] J. Sukumaran and Mark T. Holder. Dendropy: A python library for phylogenetic computing. Bioinformatics, 26:1569–1571, 2010.

[7] Lawrence Hubert and Phipps Arabie. Comparing partitions. Journal of Classification, 2(1):193–218, |mDecember 1985.

[8] Nguyen Xuan Vinh, Julien Epps, and James Bailey. Information theoretic measures for clusterings comparison: is a correction for chance necessary? In Proceedings of the 26th Annual International Conference on Machine Learning, pages 1073–1080. ACM, 2009.

[9] Marcilio CP de Souto, Ivan G. Costa, Daniel SA de Araujo, Teresa B. Ludermir, and Alexander Schliep. Clustering cancer gene expression data: a comparative study. BMC Bioinformatics, 9(1):1, 2008.

[10] Sen Song, Liang Liu, Scott V. Edwards, and Shaoyuan Wu. Resolving conflict in eutherian mammal phylogeny using phylogenomics and the multi-species coalescent model. Proceedings of the National Academy of Sciences, 109(37):14942–14947, September 2012.

[11] S. Mirarab, R. Reaz, Md S. Bayzid, T. Zimmermann, M. S. Swenson, and T. Warnow. ASTRAL: genome-scale coalescent-based species tree estimation. Bioinformatics, 30(17):i541–i548, September 2014.

